# Spontaneous Neural Activity Changes after Bariatric Surgery: a resting-state fMRI study

**DOI:** 10.1101/2021.04.01.437973

**Authors:** Yashar Zeighami, Sylvain Iceta, Mahsa Dadar, Mélissa Pelletier, Mélanie Nadeau, Laurent Biertho, Annie Lafortune, André Tchernof, Stephanie Fulton, Alan Evans, Denis Richard, Alain Dagher, Andréanne Michaud

## Abstract

1.

**Background:** Metabolic disorders associated with obesity could lead to alterations in brain structure and function. Whether these changes can be reversed after weight loss is unclear. Bariatric surgery provides a unique opportunity to address these questions because it induces marked weight loss and metabolic improvements which in turn may impact the brain in a longitudinal fashion. Previous studies found widespread changes in grey matter (GM) and white matter (WM) after bariatric surgery. However, findings regarding changes in spontaneous neural activity following surgery, as assessed with the fractional amplitude of low frequency fluctuations (fALFF) and regional homogeneity of neural activity (ReHo), are scarce and heterogenous. In this study, we used a longitudinal design to examine the changes in spontaneous neural activity after bariatric surgery (comparing pre- to post-surgery), and to determine whether these changes are related to cardiometabolic variables.

**Methods:** The study included 57 participants with severe obesity (mean BMI=43.1±4.3kg/m^2^) who underwent sleeve gastrectomy (SG), biliopancreatic diversion with duodenal switch (BPD), or Roux-en-Y gastric bypass (RYGB), scanned prior to bariatric surgery and at follow-up visits of 4 months (N=36), 12 months (N=29), and 24 months (N=14) after surgery. We examined fALFF and ReHo measures across 1022 cortical and subcortical regions (based on combined Schaeffer-Xiao parcellations) using a linear mixed effect model. Voxel-based morphometry (VBM) based on T1-weighted images was also used to measure GM density in the same regions. We also used an independent sample from the Human Connectome Project (HCP) to assess regional differences between individuals who had normal-weight (N=46) or severe obesity (N=46).

**Results:** We found a global increase in the fALFF signal with greater increase within dorsolateral prefrontal cortex, precuneus, inferior temporal gyrus, and visual cortex. This effect was more significant 4 months after surgery. The increase within dorsolateral prefrontal cortex, temporal gyrus, and visual cortex was more limited after 12 months and only present in the visual cortex after 24 months. These increases in neural activity measured by fALFF were also significantly associated with the increase in GM density following surgery. Furthermore, the increase in neural activity was significantly related to post-surgery weight loss and improvement in cardiometabolic variables, such as insulin resistance index and blood pressure. In the independent HCP sample, normal-weight participants had higher global and regional fALFF signals, mainly in dorsolateral/medial frontal cortex, precuneus and middle/inferior temporal gyrus compared to the obese participants. These BMI-related differences in fALFF were associated with the increase in fALFF 4 months post-surgery especially in regions involved in control, default mode and dorsal attention networks.

**Conclusions:** Bariatric surgery-induced weight loss and improvement in metabolic factors are associated with widespread global and regional increases in neural activity, as measured by fALFF signal. These findings alongside the higher fALFF signal in normal-weight participants compared to participants with severe obesity in an independent dataset suggest an early recovery in the neural activity signal level after the surgery.

## 2. Introduction

Obesity is a complex chronic disease (1) associated with a multiplicity of metabolic alterations, such as insulin resistance, dyslipidemia, elevated blood pressure, and chronic low-grade inflammation (2). In addition to these well-known metabolic factors, midlife obesity has also been related to subsequent cognitive impairment and is a risk factor for vascular dementia (3-5). It has been suggested that structural and functional brain alterations associated with obesity might mediate the association between obesity and cognitive and mood dysfunction (6-9).

Brain imaging studies in adults have shown associations between obesity and cortical thinning or reduction in grey matter (GM) volume in several brain regions including the orbitofrontal cortex (OFC), ventromedial prefrontal cortex, cerebellum, and temporal poles (10-13). Magnetic resonance imaging (MRI) studies have also demonstrated relationships between obesity and white matter (WM) disruption in several tracts, including the corpus callosum, internal capsule and thalamic radiation (7, 14). Several functional magnetic resonance imaging (fMRI) studies have reported differences in brain activity levels between participants who are lean and those with obesity in various resting-state networks, including the default-mode network, salience network, prefrontal and temporal lobe networks (15-19). However, the exact mechanisms underlying the links between obesity and structural and functional brain alterations remain largely unknown. Growing evidence suggests that metabolic alterations associated with obesity including chronic low-grade inflammation (20-24) and insulin resistance (25) might be of particular relevance. Using structural equation modelling in a large sample of 15,000 aging adults, a recent study found that the metabolic alterations associated with obesity might lead to cerebrovascular damage (as measured by volume of white matter hyperintensities) which could in turn affect the integrity of subcortical and cortical regions as well as cognitive and emotional functions (6).

Weight-loss interventions that induce significant metabolic and inflammatory improvements can reverse adiposity-related brain abnormalities to some extent. Bariatric surgery is an interesting model to examine the impact of sustained weight loss and long-term metabolic improvement (26-28) on brain structure and function. Our recent findings (29) confirmed the results of three studies showing an increase in WM and GM densities following bariatric surgery (30-32). These GM and WM increases were most pronounced and widespread 12 months after the surgery, and were significantly associated with post-operative weight loss and improvement of metabolic/inflammatory variables (29). To support the idea that the effects of bariatric surgery represent a resolution of adiposity-related brain abnormalities, we previously investigated whether the brain regions that show improvements after bariatric surgery are also different between individuals who are severely obese and those who are lean. Using an independent dataset from the Human Connectome Project (HCP), we found that changes in brain morphometry observed following surgery overlapped with brain differences between the obese and normal weight states, in line with the hypothesis that weight loss and/or concomitant improvement of metabolic/inflammatory alterations could be responsible for the post-operative changes in the areas previously impacted by obesity (29). Previous studies have also suggested that post-operative changes in WM and GM densities could be attributed to neural plasticity (30), recovery of brain integrity (31, 32), or changes in the composition of fiber tracts (30). However, more studies are needed to identify the underlying causes of these changes and how structural brain changes following surgery are associated with changes in brain activity (33).

Few resting-state fMRI (rsfMRI) studies have examined changes in spontaneous neural activity following bariatric surgery using various methods including the amplitude of low frequency fluctuations (ALFF), fractional-ALFF (fALFF), or regional homogeneity of neural activity (ReHo) (30, 34-36). fALFF measures the contribution of the low-frequency oscillations within a voxel relative to the entire detectable frequency range (37), and represents a marker of regional brain activity (38). ReHo measures the similarity or synchronization of the time series of a particular voxel to those of its nearest neighbors (39, 40). These signals have been previously used to detect neural activity alterations in patients with various neuropsychiatric diseases, including Alzheimer, major depressive disorder, schizophrenia, attention-deficit/hyperactivity disorder, and addiction (40, 41). Rullmann et al. were the first to use both structural and functional MRI to examine brain changes up to one year after gastric bypass surgery (RYGB) (30). They found no significant effect of the surgery on ReHo or ALFF across the whole brain, and no significant association was found between changes in ReHo or ALFF and changes in adiposity or metabolic variables. However, when restricting the analysis to significant GM clusters, they found that changes in GM density over the first year after surgery were significantly associated with elevated ReHo in the same regions. Li et al. observed increased fALFF values 4 months after sleeve gastrectomy (SG) in superior and orbitofrontal areas (36). Applying regions-of-interest analyses, a recent study found decreased ALFF in the hippocampus and increased ALFF in the posterior cingulate cortex one month following SG (35). Taken together, findings regarding changes in spontaneous neural activity following surgery are scarce and heterogeneous. Studies with longer time follow-ups are also needed to better understand the effect of bariatric surgery-induced weight loss on functional brain activity.

Thus, in the current longitudinal study, we used rsfMRI to first characterize changes in spontaneous neural activity (ReHo and fALFF) 4 months, 12 months and 24 months after bariatric surgery, comparing pre- to post-surgery changes, and to examine whether these changes are related to cardiometabolic risk factors. We also used structural MRI data to examine the effect of bariatric surgery on GM density and its association with spontaneous neural activity. We tested the hypothesis that bariatric surgery induces significant changes in spontaneous neural activity (ReHo and fALFF), and these changes are related to changes in GM density as well as improvements in cardiometabolic variables. As in our previous study (29), we used an independent dataset from HCP to test whether the change in rsfMRI signals observed following bariatric surgery is similar to that seen in individuals with normal body weight versus severe obesity.

## 3. Methods

### 3.1 Participant recruitment

Participants were recruited through the elective surgery schedule of the *Institut universitaire de cardiologie et de pneumologie de Québec*. The study sample included participants with severe obesity (mean BMI=43.1±4.3kg/m^2^) who underwent sleeve gastrectomy (SG), biliopancreatic diversion with duodenal switch (BPD), or Roux-en-Y gastric bypass (RYGB). In total, we included 57 participants at baseline (approximately 2 months prior to surgery), 36 participants at 4 months post-surgery, 29 participants at 12 months post-surgery, and 14 participants at 24 months post-surgery (**Table 1**). Inclusion criteria were the following: 1) women or men who require surgery and who meet the NIH Guidelines for bariatric surgery: BMI ≥40 kg/m^2^ or BMI≥35kg/m^2^ with comorbidities (42); 2) age between 18 and 60 years. Exclusion criteria were the following: 1) any uncontrolled medical, surgical, neurological or psychiatric condition; 2) liver cirrhosis or albumin deficiency; 3) any medication that may affect the central nervous system; 4) pregnancy; 5) substance or alcohol abuse; 6) previous gastric, oesophageal, brain or bariatric surgery; 7) gastro-intestinal inflammatory diseases or gastro-intestinal ulcers; 8) severe food allergy; and 9) contraindications to MRI. The study was approved by the Research Ethics Committee of the *Centre de recherche de l’Institut universitaire de cardiologie et pneumologie de Québec (approval number 2016-2569)*. All participants provided written informed consent to participate in the study.

**Table 1.** Characteristics of participants at baseline, 4, 12 and 24 months after bariatric surgery.

| Variable | Baseline | 4 months | 12 months | 24 months | *p* value |
| --- | --- | --- | --- | --- | --- |
| *N* | 57 | 36 | 29 | 14 | - |
| Sex (F : M) | 42 : 15 | 26 : 10 | 24 : 5 | 9 : 5 | 0.591 |
| Surgery type (SG : RYGB : BPD) | - | 23 : 7 : 6 | 17 : 7 : 5 | 8 : 4 : 2 | 0.443 |
| Type 2 Diabetes Mellitus (Y : N) | 16:41 | - | - | - | - |
| Age (years) | $45.8 \pm 8.5$ | $47.2 \pm 8.1$ | $49.0 \pm 7.6$ | $48.6 \pm 8.7$ | 0.325 |
| Weight (kg) | $121.0 \pm 14.0$ | $93.8 \pm 10.3$ | $80.1 \pm 10.9$ | $82.5 \pm 14.5$ | **< 0.001** |
| BMI (kg/m<sup>2</sup>) | $43.1 \pm 4.3$ | $34.2 \pm 3.3$ | $29.3 \pm 3.8$ | $28.8 \pm 4.3$ | **< 0.001** |
| Waist circumference (cm) | $129 \pm 11$ | $111 \pm 11$ | $100 \pm 11$ | $100 \pm 13$ | **< 0.001** |
| Neck circumference (cm) | $42 \pm 3$ | $37 \pm 3$ | $35 \pm 3$ | $36 \pm 3$ | **< 0.001** |
| Excess weight loss (%) | - | $47.0 \pm 9.2$ | $70.4 \pm 14.9$ | $72.2 \pm 17.5$ | **< 0.001** |
| Total weight loss (%) | - | $21.9 \pm 4.0$ | $32.6 \pm 6.7$ | $33.2 \pm 8.7$ | **< 0.001** |
| Systolic blood pressure (mmHg) | $135 \pm 17$ | $122 \pm 15$ | $122 \pm 18$ | $117 \pm 12$ | **< 0.001** |
| Diastolic blood pressure (mmHg) | $80 \pm 11$ | $73 \pm 12$ | $73 \pm 12$ | $70 \pm 12$ | **0.003** |
| Fasting glycemia (mmol/L) | $6.5 \pm 1.7$ | $5.2 \pm 0.9$ | $4.8 \pm 0.9$ | $5.0 \pm 0.7$ | **< 0.001** |
| HbA1c (%) | $5.8 \pm 0.9$ | $5.3 \pm 0.6$ | $5.1 \pm 0.4$ | $5.1 \pm 0.4$ | **< 0.001** |
| Fasting insulin (pmol/L) | $170.3 \pm 96.7$ | $69.7 \pm 37.1$ | $45.1 \pm 30.4$ | $46.3 \pm 22.5$ | **< 0.001** |
| HOMA-IR index | $7.3 \pm 4.8$ | $2.3 \pm 1.5$ | $1.4 \pm 1.6$ | $1.5 \pm 0.9$ | **< 0.001** |
| Total chol./HDL-chol. (mmol/L) | $3.9 \pm 1.0$ | $3.5 \pm 1.0$ | $2.9 \pm 0.8$ | $2.8 \pm 0.7$ | **< 0.001** |
| LDL-cholesterol (mmol/L) | $2.8 \pm 1.1$ | $2.4 \pm 1.0$ | $2.3 \pm 0.7$ | $2.2 \pm 0.8$ | 0.059 |
| HDL-cholesterol (mmol/L) | $1.2 \pm 0.3$ | $1.2 \pm 0.3$ | $1.5 \pm 0.3$ | $1.6 \pm 0.2$ | **< 0.001** |
| Triglycerides (mmol/L) | $1.8 \pm 0.9$ | $1.5 \pm 1.0$ | $1.1 \pm 0.5$ | $1.1 \pm 0.6$ | **0.003** |

Results are presented as mean ± SD; repeated-measures ANOVA comparing baseline, 4 months 12 months and 24 months post-surgery sessions. SG, sleeve gastrectomy; RYGB, Roux-en-Y gastric bypass; BPD, biliopancreatic derivation with duodenal switch; BMI, body mass index; HbA1c, hemoglobin A1c or glycated hemoglobin; HOMA-IR, homeostatic model assessment for insulin resistance; HDL, high-density lipoprotein; LDL, low-density lipoprotein, chol., cholesterol

### 3.2 Surgical procedures

Laparoscopic SG, a restrictive surgery consisting of 150 to 250 cm^3^ vertical gastrectomy on a 34 French bougie starting 4 to 5cm proximal to the pylorus (43), was performed in most of the participants (n=23). Laparoscopic BPD, a mixed-surgery combining restrictive and malabsorptive mechanisms by creating a 150-250 cm^3^ vertical SG on a 34 French bougie and duodeno-ileal anastomosis 250 cm from ileocecal valve (including a common limb of 100-cm) (44), was performed in 6 participants. Laparoscopic RYGB, another mixed-surgery in which a proximal gastric pouch of 30-50cm^3^ is created and anastomosed to the proximal small bowel by bypassing the first 100 cm and bringing a 100 cm alimentary limb on the gastric pouch, was performed in 7 participants.

### 3.3 Study design and experimental procedure

The study design has been described in detail in (29). Briefly, participants were assessed 2 months prior to as well as 4, 12, and 24 months post-surgery. Before each MRI session, participants underwent a physical examination, including blood pressure and anthropometric measurements: body weight and body composition using calibrated bioelectrical impedance scale (InBody520, Biospace, Los Angeles, CA, or Tanita DC-430U, Arlington Heights, IL) as well as height, waist circumference, and neck circumference using standardized procedures. We calculated BMI (kg/m^2^), percentage of excess weight loss and percentage of total weight loss as previously described (29). Fasting blood biochemistry was also obtained at each visit. Plasma levels of cholesterol, high-density lipoproteins, low-density lipoproteins, triglycerides, glycated hemoglobin (HbA1c), glucose and insulin were measured. The homeostasis model assessment insulin resistance (HOMA-IR) index was calculated using this formula: (fasting glucose (mmol/L) x fasting insulin (mU/L))/22.5.

### 3.4 MRI acquisition

MRI acquisition was performed on the morning of each visit (starting between 9:00 and 10:30). Participants were asked to fast 12h before the MRI session and received a standardized beverage meal (Boost original, Nestle Health Science) 1h before the MRI session to control for hunger (29). MRI scans were performed on a 3T whole-body MRI scanner (Philips, Ingenia, Philips Medical Systems) equipped with a 32-channel head coil at the *Centre de recherche de l’Institut universitaire de cardiologie et pneumologie de Québec*. Participants were placed in the scanner in supine position. At each visit, the MRI protocol included anatomical T1-weighted three-dimensional (3D) turbo field echo images, rsfMRI and a task state fMRI for food-cue reactivity. In the current study, only results from T1-weighted anatomical images and rsfMRI are presented. The following parameters were used for the T1-weighted images: 176 sagittal 1.0 mm slices, repetition time/echo time (TR/TE) = 8.1/3.7 ms, field of view (FOV) = 240 x 240 mm^2^, and voxel size = 1 x 1 x 1 mm. During the rsfMRI, participants were instructed to fixate on a cross in the middle of a black screen and relax, but not fall asleep. T2*-weighted echo planar images (EPI) with blood-oxygen-level-dependent contrast were acquired for ∼10 minutes. Each volume comprised 45 transverse slices, a TE = 30 ms, a TR = 2750 ms, a voxel size of 3 x 3 x 3 mm, a FOV = 240 x 240 mm^2^, and a flip angle of 80°.

#### 3.4.1 Resting-state fMRI processing

T1-weighted MRI and rsfMRI images were converted to BIDs format (45). All subjects were then preprocessed using fMRIPrep pipeline version v1.3.0.post2 (46). Briefly, T1-weighted MRIs were corrected for intensity non-uniformity, skull-stripped, and registered linearly and nonlinearly to MNI-152-2009c template (47). The T1-weighted MRIs were then segmented into cerebrospinal fluid (CSF), WM, and GM tissues. rsfMRIs slices were first coregistered to the mean rsfMRI image, motion parameters were estimated. The rsfMRI image was then coregistered to the participants’ native T1-weighted image and then native T1-weighted to MNI template registration was applied to each participant’s rsfMRI image. Signals from CSF, WM, and GM were extracted from the rsfMRI time series.

We used the output of fMRIPrep pipeline and further applied SPM12 (https://www.fil.ion.ucl.ac.uk/spm/software/download/) (48) slice timing, detrending for fMRI time series fMRI image (using LMGS method) as well as correcting the time series for motion parameters and average CSF and WM signal. Finally, we used REsting State fMRI data analysis Toolkit (REST version v1.8; https://www.nitrc.org/projects/rest/) (49) to calculate ReHo and fALFF using 0.01 and 0.08 as low and high cut off frequencies respectively.

#### 3.4.2 Voxel-based morphometry measurements

As in our previous study (29), GM densities were measured from T1-weighted anatomical MRIs of each participant using a standard voxel-based morphometry (VBM) pipeline (10). The preprocessing steps included: 1) image denoising (50); 2) intensity non-uniformity correction (51); and 3) image intensity normalization into range (0-100) using histogram matching. Images were then first linearly (using a nine-parameter registration) and then nonlinearly registered to MNI-152-2009c template as part of the ANIMAL software (52) and segmented into GM, WM and CSF images. These steps remove global differences in the size and shape of individual brains and transform individual GM density maps to the standardized MNI-152-2009c template space. VBM analysis was performed using MNI MINC tools (http://www.bic.mni.mcgill.ca/ServicesSoftware/MINC) to generate GM density maps representing voxel-wise GM concentration.

### 3.5 Parcellation

Schaffer functional MRI parcellation at 1000 regions (https://github.com/ThomasYeoLab/CBIG/tree/master/stable_projects/brain_parcellation/Schaefer2018_LocalGlobal)(53) was used to extract regional information for all measurements of interest. Schaeffer-1000 regions are derived based on the Yeo et al. (54) seven large scale functional networks, and subdivides each network to multiple regions based on the functional similarity within regions. For subcortical regions, we used the atlas developed by Xiao et al. (55) including 11 subcortical regions in each hemisphere (http://nist.mni.mcgill.ca/?p=1209) (i.e., amygdala, hippocampus, thalamus, caudate, nucleus accumbens, putamen, globus pallidus internal/external, red nucleus, substantia nigra, and subthalamic nucleus). All statistical analyses performed are based on the data extracted from 1022 regions of the combined Schaeffer-Xiao parcellation.

### 3.6 Statistical analyses

We used repeated-measures ANOVA to compare the clinical characteristics of participants at baseline, 4, 12, and 24 months after bariatric surgery. We also used linear-mixed effects models to assess the effect of surgery on metabolic variables as well as ReHo and fALFF signals across the cortical and subcortical brain regions. MRI sessions were used as the fixed effect of interest (4-month, 12-month, and 24-month visits contrasted against baseline scan before surgery) and subjects were used as the random effect. All models included age, sex, BMI, and type 2 diabetes status at baseline, as well as surgery type (SG, BPD and RYGB) as covariates. A similar model was used to investigate the relationship between metabolic variables and fALLF signal. All results were corrected for multiple comparisons using Benjamini-Hochberg false discovery rate (BH-FDR) (56) correction with q=0.05.

To examine the impact of the surgery on the relationship between changes in fALFF signal and changes in GM density (as previously measured in (29)), we used a linear-mixed effects model to estimate the effect of surgery on each signal (i.e. fALLF or GM) as described in the previous paragraph. The association analysis between the resulting statistical estimates of the changes of GM density and fALFF were performed using Spearman correlation for 4, 12, 24 months after surgery separately. Spin-test analysis with N=10,000 permutations was used to evaluate the degree of similarity between surgery-related fALFF and GM density changes. Spin-test is a permutation-based method developed to evaluate the spatial correspondence between maps of the human brain while accounting for spatial autocorrelation (using in https://github.com/frantisekvasa/rotate_parcellation) (57, 58). Spin-test results were also corrected for multiple testing using Bonferroni correction. All statistical analyses were performed using MATLAB R2018a.

### 3.7 Validation in an independent dataset from the Human Connectome Project

We used rsfMRI data from HCP as an independent dataset for validation. All participants with BMI higher than 35 kg/m^2^ (N=46) were included. These participants were individually matched for age, sex and ethnicity with a group of unrelated normal weight participants from the same HCP dataset (N=46). For all participants, FIX-preprocessed rsfMRI data nonlinearly registered to the MNI space was downloaded from the data repository (https://db.humanconnectome.org). Briefly, this preprocessing uses independent component analysis (ICA) to detect motion and other artifacts. These ICA components are then regressed out from the time series as explained in Salimi-Khorshidi et al. (59). We included both left-right and right-left sequences from the first session (except for one participant with missing scans for which we included data from the second session). fALFF signal was calculated was for the bariatric surgery sample. The fALFF results from left-right and right-left scans were averaged, and the average was used as the measure of interest. All results in the validation study are based on Student’s paired t-tests analysis between the severely obese and normal-weight subgroups. For consistency, we translated the t-values to the regional group difference estimates (as derived from the linear models) for visualization purposes. Since the subgroups are matched on age and sex and there was no intervention in the HCP group (i.e. surgery), no covariate was used for this analysis.

## 4. Results

### 4.1 Clinical characteristics of participants

**Table 1** shows the clinical characteristics of the participants included in this study. At baseline, the study sample included 42 women and 15 men. As expected, patients at baseline demonstrated significant metabolic anomalies including insulin resistance and dyslipidemia. However, only 16 of 57 patients had a diagnosis of Type 2 diabetes mellitus. The most frequently performed surgery was SG. The mean total weight loss was 21.9% ± 4.0 after 4 months, 32.6% ± 6.7 after 12 months, and 33.2% ± 8.7 after 24 months. Significant improvements in metabolic variables (blood pressure, glucose homeostasis, and lipid profile) were observed after surgery.

### 4.2 Effect of bariatric surgery on global fALFF and ReHo signals

There was a significant increase in global fALFF signal after bariatric surgery compared to baseline. The increase was higher at 4 months post-surgery (t = 3.99, p < 0.001). Even though fALFF signal was still significantly higher than baseline after 12 months (t = 2.47, p = 0.016) and 24 months (t = 2.22, p = 0.037), these increases were less pronounced compared to the 4-month visit. This effect was seen in cortical and subcortical areas, but subcortical areas showed a lower increase compared to cortical areas. We did not observe significant effect for other variables included in the model, such as age, sex, BMI at baseline and type 2 diabetes status at baseline, as well as surgery type. No significant change was observed in ReHo signal at the global GM level (p > 0.05, data not shown).

### 4.3 Effect of bariatric surgery on fALFF and ReHo signal across brain regions

We further examined the change in fALFF and ReHo signals across brain areas using the 1022 regions of the combined Schaeffer-Xiao parcellation. **Figure 1 A-C** shows the regional increase in fALFF signal for 4, 12, and 24 months after surgery, respectively (after FDR correction for multiple comparison). While the short-term increase in fALFF was widespread and present across the brain, the increase was greater in the dorsolateral prefrontal cortex, visual cortex, followed by anterior cingulate gyrus, precuneus, middle and inferior temporal gyrus. However, only the visual cortex showed a significant increase in fALFF signal 24 months after surgery. We repeated the same analysis for mean ReHo and found no significant effect of surgery over time after correction for multiple comparisons.

**Figure 1.**
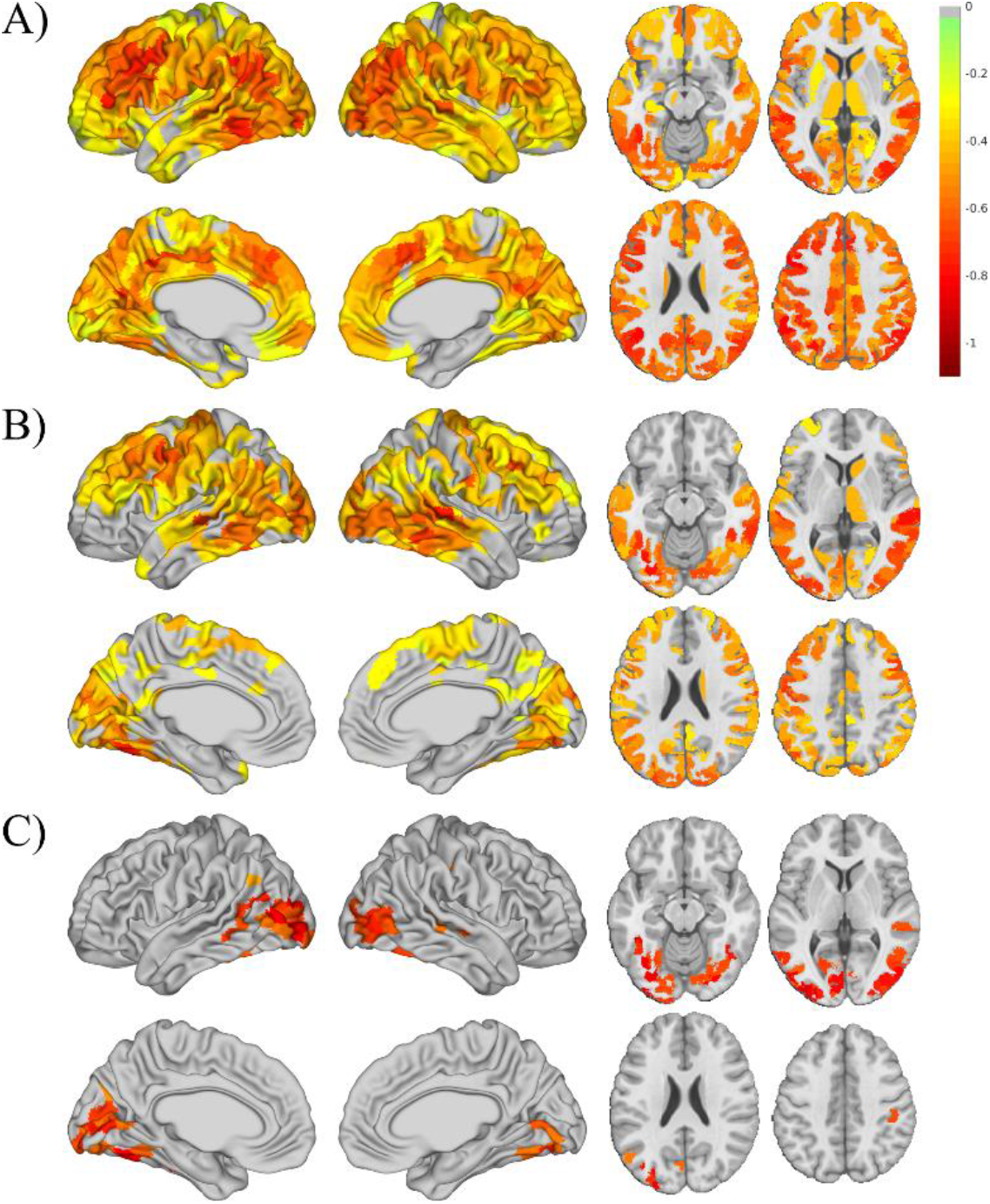
The regional changes of fALFF signal **A)** 4 months, **B)** 12 months, and **C)** 24 months after bariatric surgery. Visualization in axial sections are shown for z = −14, 4, 23, and 42 in MNI space.

We further mapped the increase in fALFF signal to the 7 functional networks (Yeo et al. 2011(54)) underlying the Schaeffer 1000 cortical regions (**Figure 2**). All networks showed an initial increase followed by a drop which was still higher than the baseline level after 24 months (i.e. positive estimate values from mixed-effect model). Dorsal attention, control, and default mode networks showed greater increase initially. However, the increases in visual and somatomotor networks were more stable over time. The limbic network showed the lowest increase in signal compared to baseline.

**Figure 2.**
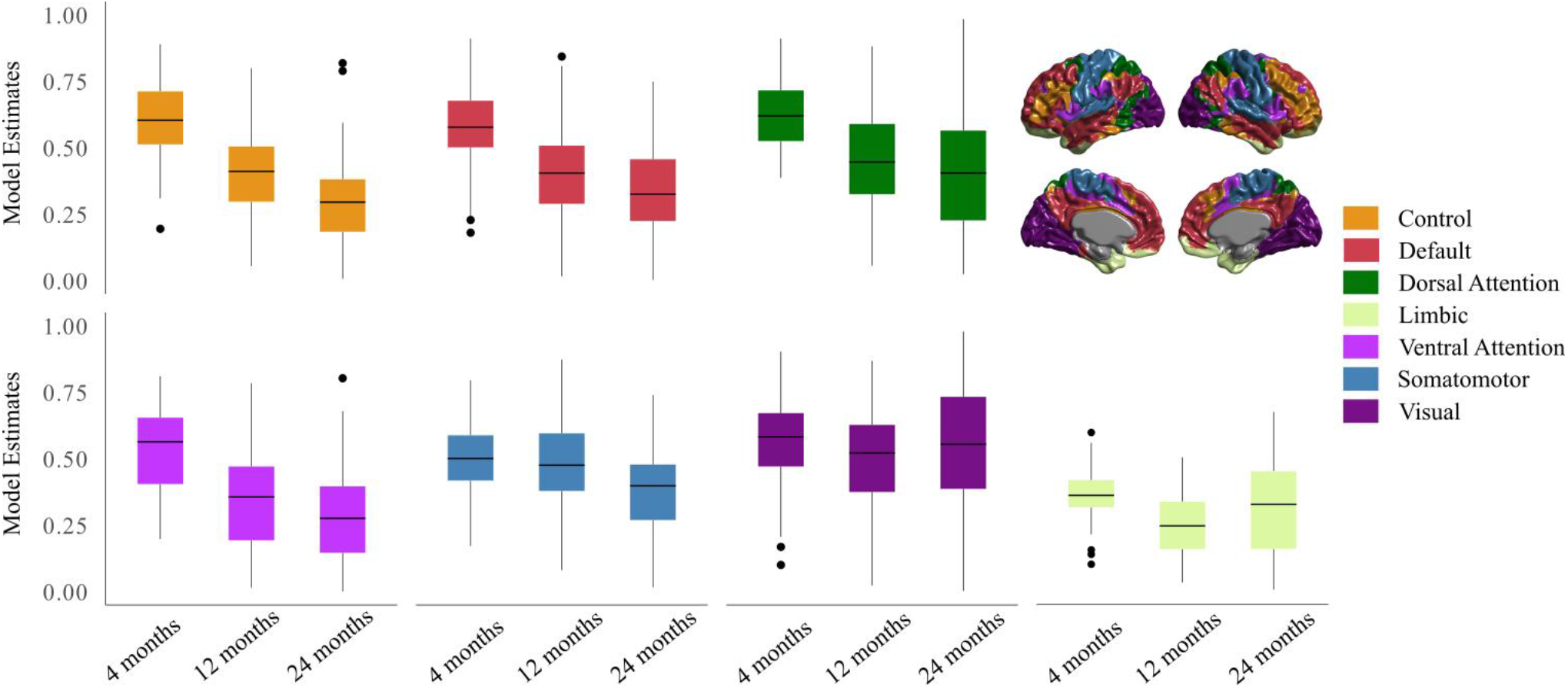
fALFF signal changes after bariatric surgery within each of the 7 large scale functional networks from Yeo et al. 2011 (54), based on the estimate values from linear mixed-effects models.

### 4.4 Effect of bariatric surgery on GM density and its association with fALFF signal

We then examined the association between changes in GM density and in fALFF signal after surgery. We used the estimated effect sizes resulting from the mixed-effects models to examine the effect of surgery on GM density (similar to the fALFF signal). We then examined the association between the changes in GM and fALFF as reflected by these estimates. We found a significant correlation between the effect of surgery on fALFF and GM density after 4 months (ρ=0.23, p-spin-test = 0.0005), 12 months (ρ=0.42, p-spin-test = 0.0000), and 24 months (ρ=0.27, p-spin-test = 0.0000), suggesting that the increase in neural activity measured by fALFF is positively associated with the increase in GM density following surgery (**Figure 3**). We further calculated the correlation between the GM and fALFF within each of the 7 functional networks for each visit (**Table 2**). There was no significant association between the change in GM and fALFF in limbic and salient ventral attention networks, while other networks showed positive and significant association between the two measurements.

**Table 2.** The association between change in fALFF signal and GM density after surgery within seven large scale brain functional networks. Results are based on Spearman rank correlation and p-values are corrected for multiple comparisons using Bonferroni method.

| Brain functional networks | 4 month |  | 12 month |  | 24 month |  |
| --- | --- | --- | --- | --- | --- | --- |
| | $\rho$ | p-value | $\rho$ | p-value | $\rho$ | p-value |
| Control | 0.06 | 1 | **0.54** | **<0.001** | **0.29** | **0.015** |
| Default Mode | **0.27** | **0.001** | **0.47** | **<0.001** | **0.23** | **0.010** |
| Dorsal Attention | 0.21 | 0.43 | **0.48** | **<0.001** | **0.51** | **<0.001** |
| Limbic | 0.20 | 1 | 0.12 | 1 | 0.16 | 1 |
| Ventral Attention | 0.20 | 0.67 | 0.24 | 0.15 | 0.04 | 1 |
| Somatomotor | **0.28** | **0.001** | **0.40** | **<0.001** | **0.36** | **<0.001** |
| Visual | **0.30** | **0.001** | **0.38** | **<0.001** | **0.38** | **<0.001** |

**Figure 3.**
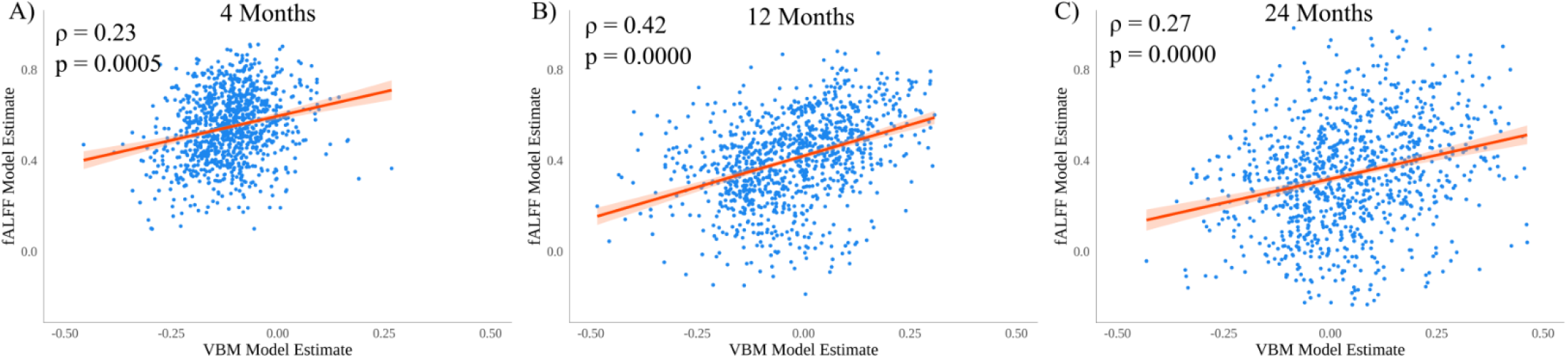
The association between change in fALFF signal and GM **A**) 4 months, **B**) 12 months and **C**) 24 months after bariatric surgery. The values are based on Spearman rank correlation and the p-values are measured using permutation based on the spin test.

### 4.5 Associations between changes in fALFF signal and adiposity/metabolic variables following bariatric surgery

We next used mixed-effects modeling analysis to examine the associations between regional changes in fALFF signal and adiposity/metabolic variables after bariatric surgery, accounting for age, sex, BMI, and type 2 diabetes status at baseline, as well as surgery type. **Figure 4** shows the significant regional changes of fALFF signal in association with adiposity or metabolic variables. Significant associations were observed between post-operative reduction in BMI and increased fALFF signal in several brain regions, mainly in temporal-occipital cortex, lingual gyrus, precuneus, postcentral and precentral gyrus, orbitofrontal cortex, cingulate gyrus and caudate (**Figure 4A**). Similar results were observed with post-operative reduction in waist circumference or percentage of excess or total weight loss (data not shown). Reduced HOMA-IR index following surgery was also significantly associated with increased fALFF signal in several brain regions, mainly in temporal-occipital regions, middle frontal regions, cingulate gyrus, postcentral and precentral gyrus, caudate and thalamus (**Figure 4B**). Similar results were observed with reduced plasma insulin levels. Significant associations were also observed between reduced diastolic blood pressure post-surgery and increased fALFF signal (**Figure 4C**). These associations were found in the same brain regions, but were more restricted (mainly in temporal-occipital regions, precentral and postcentral gyrus, precuneus, superior frontal gyrus and frontal operculum cortex). Similar results, but to a lesser extent were found with improvements in systolic blood pressure (data not shown). No significant association was observed after FDR correction between fALFF signal and plasmatic levels of glucose, HbA1c, LDL-cholesterol, HDL-cholesterol, and triglycerides (data not shown).

**Figure 4.**
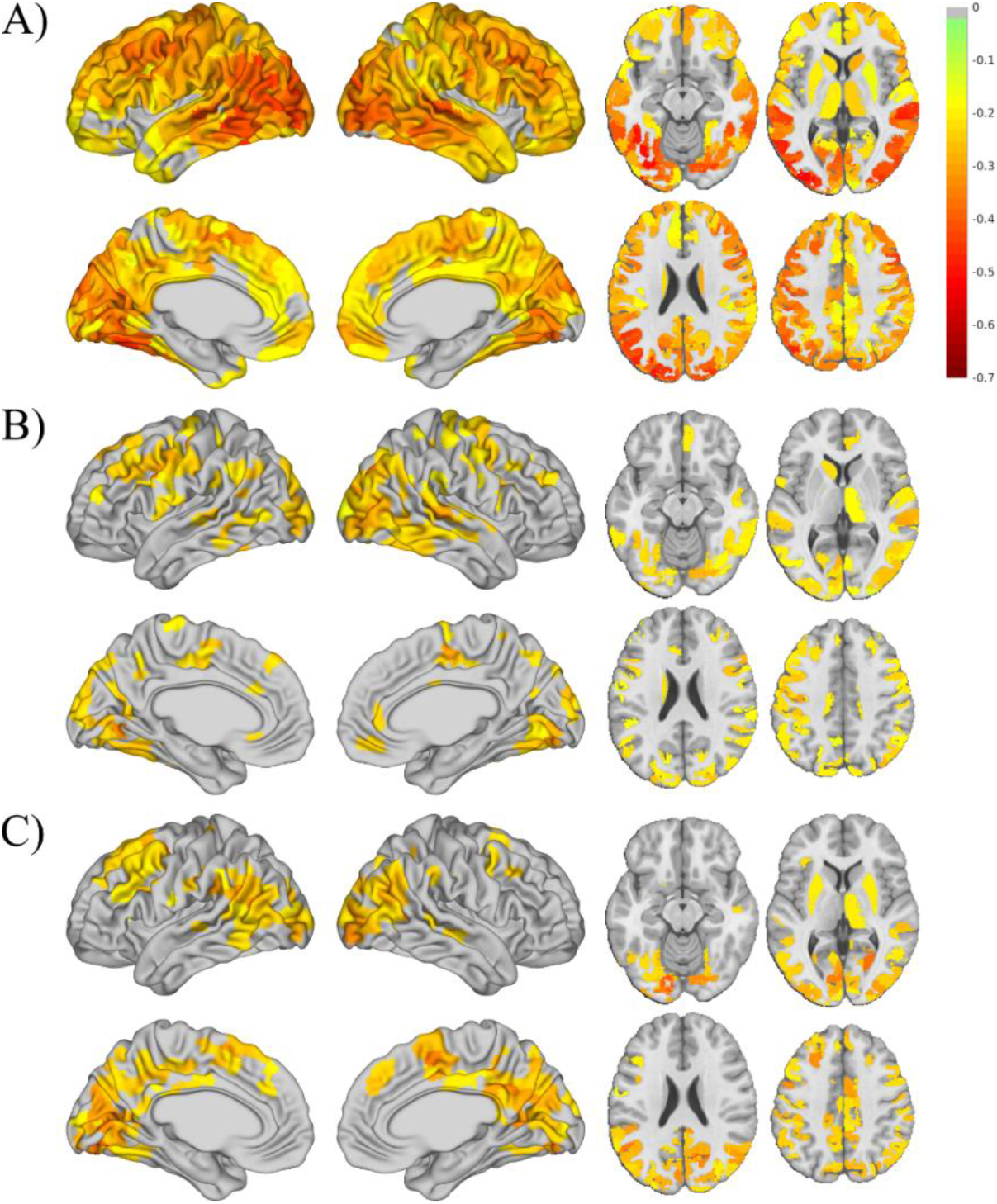
The regional changes of fALFF signal in association with **A)** BMI, **B)** HOMA-IR, and **C)** diastolic blood pressure during a 24 months period after bariatric surgery. Visualization in axial sections are shown for z = −14, 4, 23, and 42 in MNI space. BMI: Body mass index; HOMA-IR: homeostasis model assessment insulin resistance index

### 4.6 Effect of obesity on fALFF signal: Validation in an independent dataset

Similar to our validation analysis in (29), we used the rsfMRI from HCP to test whether the increase in fALFF signals observed following surgery is associated with BMI-related differences in an independent dataset. We found that participants with normal weight have a higher global fALFF signal compared to those with severe obesity (t = 2.65, p < 0.01). We also found significantly higher fALFF signal in normal-weight participants compared to those with severe obesity, mainly in dorsolateral and dorsomedial frontal cortex, precuneus, as well as middle and inferior temporal gyrus (**Figure 5A**). The regional pattern of the higher fALFF signal in participants who had normal body weight compared to those with obesity correlated significantly with the increased fALFF signal after 4 months of surgery compared to baseline (ρ= 0.237, p < 0.0001, p-spin-test= 0.0000, **Figure 5B**). Default mode, ventral and dorsal attention and control networks showed the strongest associations, while the associations were less strong but still significant within somatomotor, visual, and limbic networks. However, we found no significant relationship between the higher fALFF pattern in the control group in HCP and the increase of fALFF signal 12 months (ρ= 0.026, p= 0.39), or 24 months (ρ= −0.009, p=0.77) after surgery.

**Figure 5.**
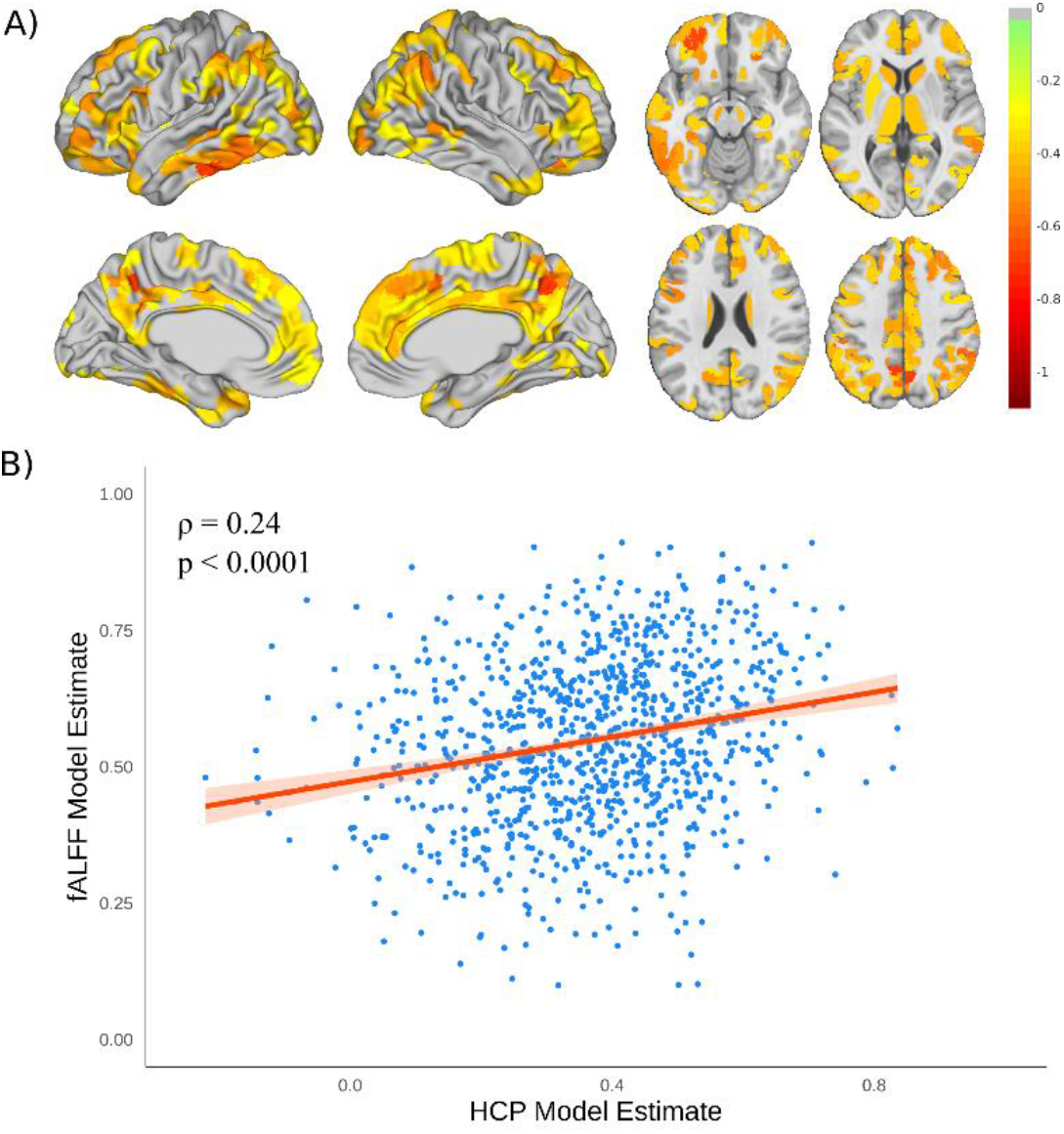
fALFF signal results from HCP dataset. **A)** The pattern of higher fALFF signal in participants who had normal body weight compared to those with severe obesity within HCP dataset (results are shown after FDR correction). Visualization in axial sections are shown for z = −14, 4, 23, and 42 in MNI space. **B)** The association between change in fALFF signal 4 months compared to baseline and the effect size of higher fALFF signal in normal-weight participants in HCP dataset.

## 5. Discussion

Our study provides evidence that bariatric surgery-induced weight loss and concomitant metabolic improvement are associated with widespread global and regional increases in resting neural activity, as indexed by fALFF. The increase in fALFF was greater 4 months post-surgery principally in prefrontal, precuneus, occipital and middle/inferior temporal regions. Interestingly, the increase in neural activity measured by fALFF was strongly associated with the increase in GM density following surgery. Furthermore, the increase in fALFF was associated with post-surgery reduced adiposity and improvements in cardiometabolic health. Using an independent dataset from HCP, we found that normal-weight participants had higher global and regional fALFF signal, in a pattern that spatially overlapped with the post-surgery changes. The effects were especially prominent in heteromodal cortex including the control, default mode, and dorsal attention networks. On the other hand, another measure of resting neuronal activity, ReHo showed no significant effect of bariatric surgery. Our findings support the idea that bariatric surgery may lead to a rapid resolution of adiposity-related brain abnormalities, along with widespread improvements in metabolism.

fALFF and ReHo have been used in other populations targeting individuals with mild cognitive impairment as well as patients with various neuropsychiatric diseases (e.g. major depressive disorder, schizophrenia, attention-deficit/hyperactivity disorder and addiction) and neurodegenerative diseases (e.g. Alzheimer’s disease and frontotemporal dementia) to detect neural activity (40, 41, 60, 61). However, very few studies have previously examined the effect of bariatric surgery on spontaneous neural activity as assessed with fALFF or ReHo (30, 34-36). We found widespread increase in fALFF 4 months after surgery, with greater increases in dorsolateral prefrontal cortex, precuneus, occipital as well as middle and inferior temporal regions. Interestingly, the increase in neural activity became more regionally limited over time, with only the visual cortex showing a significant increase in fALFF 24 months after surgery compared to baseline. These results are consistent with a previous study from Li et al. showing increased neural activity, as measured with fALFF, in the orbitofrontal cortex, superior frontal gyrus and gyrus rectus 4 months after sleeve gastrectomy (36). Using ROI analysis, Zhang et al. also found increased neural activity as assessed with ALFF in the posterior cingulate cortex 1 month after sleeve gastrectomy (35). Considering that neural activity was reduced in the orbito/superior frontal regions in participants with obesity, Li et al. suggested that bariatric surgery induces recovery of brain anomalies present in obesity (36). Using an independent dataset from HCP, we also found that the post-operative changes in fALFF signal at 4 months were in brain regions that showed alterations in obesity, especially regions of the prefrontal cortex known to be implicated in cognitive control (33, 62). Considering that the increase in neural activity in brain regions involved in control network is greater at short-term (4 months versus 12 and 24 months), it could explain why we found no significant association between the increase in fALFF signal at 12- or 24-months and BMI-related differences in fALFF signal in HCP dataset. Taken together, our results suggest an early-stage reversal toward a normal neural activity pattern following bariatric surgery.

We further found that the increase in neural activity measured by fALFF was associated with the increase in GM density following bariatric surgery, which suggest a regional correspondence between the two measurements and a global effect of bariatric surgery on brain status. As previously mentioned, the changes in fALFF were more widespread and pronounced after 4 months, while the changes in GM density were more pronounced 12 months after bariatric surgery (29), which suggests that morphological brain changes after surgery might take more time to occur than functional changes. As our current sample lacks sufficient participant data at all follow-ups, it was not possible to establish a causal relationship between functional and morphological changes. The higher VBM GM changes observed after 12 months (29) might also explain why we observed a stronger association between neural activity and GM density at 12 months.

While Rullman et al. failed to observe significant changes in ALFF or ReHo values 6 and 12 months following RYGB compared to baseline, they did find significant relationships between elevated ReHo after surgery and GM density changes in cortical and subcortical regions (30). However, they found no significant associations between ALFF and GM density. Their results differ from our study mainly due to methodological differences between studies, such as the analyses used to measure spontaneous neural activity (ALFF versus fALFF), sample sizes, the type of surgery and the timeline. While ALFF and fALFF are closely related conceptually, the normalization factor included in fALFF could potentially decrease the variance in the signal and consequently increase the statistical power. Furthermore, the normalized nature of fALFF signal makes it easier to interpret the findings between groups and across brain regions. For example, while non-specific to neural activity, a general increase in signal due to surgery can be detected in ALFF as a significant change, whereas an increase in fALFF will only be observed if the fraction related to neural activity shows a significant increase. Therefore, fALFF findings are less likely to be prone to the general non-neural signal changes. More studies with larger sample sizes are needed to further confirm these findings.

The increase in neural activity after bariatric surgery was correlated with the degree of weight loss and concomitant improvement in cardiometabolic variables. More specifically, we found significant associations between the increase in fALFF and post-operative improvement in insulin resistance (as measured by HOMA-IR index) and blood pressure. No significant association was observed with changes in lipid profile or levels of fasting glucose/HbA1c. Rullmann et al. found that markers of neural activity prior to surgery were not associated with changes in BMI, lipid profile, HbA1c, TSH or CRP after 6 months or 12 months post-surgery (30). While we cannot establish the specific contribution of each metabolic parameter or make any causal inference in our study, our findings support the idea that the increase in neural activity and GM volume could be a consequence of a better blood circulation and improved insulin sensitivity following weight-loss surgery. Indeed, there is a rapid improvement in artery endothelial function following bariatric surgery (63, 64), which is also associated with the remission of hypertension (65) and reduction of atherosclerosis (66). The observed improvement in cardiometabolic/vascular function may well connect to angiogenesis, improved integrity of the neurovascular unit and reduced brain gliosis that could promote dendritic branching and synaptogenesis and influence MRI measurements (67, 68). Changes in cerebral blood flow and oxygen concentration observed after the surgery could also influence a portion of the MR-based structural biomarkers, such as GM density and cortical thickness (69). The improvement in insulin sensitivity following surgery could also be an important mechanism to explain the changes in neural activity since brain insulin resistance has previously been associated with alterations in cerebral glucose metabolism, brain atrophy, and cognitive alterations (70, 71). Other mechanisms have been proposed to explain the effects of bariatric surgery on brain structure and function, including the improvement of chronic low-grade inflammation, changes in incretin/gut peptide secretion (35, 72) and microbiota products that impact the brain (33, 73, 74). A better understanding of the cellular/biological mechanisms through which the cardiometabolic changes following bariatric surgery influence human brain function and structure is of particular importance.

Our study is not without limitations. We did not have data from all the visits for every participant included in the study. This is a significant limitation, because some brain changes showed greater effects at short term (4 months) compared to long term. Furthermore, we used the participants as their own control (comparing pre-op versus post-surgery), because it is difficult to include a control group (without surgery) with the same magnitude of weight loss. To support the idea that the post-operative brain changes go in the direction of a resolution of brain abnormalities related to obesity, we used rsfMRI data from participants from the HCP who had normal body weight or severe obesity as an independent dataset for validation. However, this population was younger and had less metabolic comorbidities compared to our bariatric participants. Furthermore, some rsfMRI parameters were different in HCP compared to our study, with HCP having a higher spatial and temporal resolution (TR=700ms, a voxel size of 2 x 2 x 2 mm^3^) which can result in different signal-to-noise ratios and statistical power.

In conclusion, bariatric surgery-induced weight loss is associated with widespread global and regional increases in neural activity, particularly in the short term. This increase in neural activity is related to the post-operative increase in GM density and the improvement in cardiometabolic health. These early post-operative changes were associated with functional brain differences between participants with normal weight and those with severe obesity, supporting the idea that the early effect of bariatric surgery on neural activity could be a resolution of an adiposity-related brain alteration.

## 6. Acknowledgments

We would like to acknowledge the contribution of surgeons, nurses, the medical team of the bariatric surgery program at IUCPQ, MRI technicians, Xavier Moreel, Coordinator of the Plateforme d’imagerie avancée at IUCPQ, and Guillaume Gilbert, Engineer, Phillips as well as the collaboration of participants.

